# Deletion of Sphingosine 1-Phosphate Receptor 1 in cardiomyocytes during development leads to abnormal ventricular conduction and fibrosis

**DOI:** 10.1101/2020.03.15.993105

**Authors:** Ryan Jorgensen, Meghna Katta, Jayne Wolfe, Desiree F. Leach, Jerold Chun, Lisa D. Wilsbacher

**Author notes:** Corresponding author: Lisa D. Wilsbacher, Northwestern University Feinberg School of Medicine, Simpson Querrey Biomedical Research Center 8-404, 303 E. Superior St., Chicago, IL 60611.

## Abstract

Sphingosine 1-phosphate receptor 1 (S1P_1_, encoded by *S1pr1*) is a G protein-coupled receptor that signals in multiple cell types including endothelial cells and cardiomyocytes. Cardiomyocyte-specific deletion of *S1pr1* during development leads to ventricular noncompaction, with one-third of mutant mice surviving to adulthood. Adult survivors of embryonic cardiomyocyte *S1pr1* deletion showed cardiac hypertrabeculation consistent with ventricular noncompaction. Surprisingly, systolic function in mutant mice was preserved through at least one year of age. Cardiac conduction was abnormal in cardiomyocyte *S1pr1* mutant mice, with prolonged QRS intervals in mutants as compared with littermate control mice. Immunostaining of hearts from *S1pr1* mutant embryos displayed a zone of intermediate Connexin 40 expression in the trabecular myocardium. However, we observed no significant differences in Connexin 40 and Connexin 43 immunostaining in hearts from adult survivors of embryonic cardiomyocyte *S1pr1* deletion, which suggests normalized development of the ventricular conduction system in mutant mice. By contrast, the adult survivors of embryonic cardiomyocyte *S1pr1* deletion showed increased cardiac fibrosis as compared with littermate controls. These results demonstrate that ventricular hypertrabeculation caused by embryonic deletion of cardiomyocyte *S1pr1* leads to cardiac fibrosis, which contributes to abnormal ventricular conduction. These results also reveal conduction abnormalities in the setting of hypertrabeculation with normal systolic function, which may be of clinical relevance in humans with ventricular hypertrabeculation.

## INTRODUCTION

Sphingosine-1-phosphate (S1P) is a bioactive lipid that acts via one of five G protein-coupled receptors (GPCRs) named S1P_1_ - S1P_5_ [1, 2]. *S1pr1*, the gene that encodes S1P_1_, is highly expressed in endothelial cells, lymphocytes, neuronal cells, and cardiomyocytes [3-6]. Global and endothelial-specific knockout of *S1pr1* in mice causes abnormal vessel development and death at midgestation due to defective sprouting angiogenesis [3, 7-9]. We showed that cardiomyocyte-specific loss of *S1pr1* during development led to ventricular septal defects and ventricular noncompaction [6]. These structural abnormalities caused perinatal death in 68% of mutant mice; however, the remaining *S1pr1* cardiomyocyte mutant mice survived to adulthood [6].

The ventricular conduction system (VCS) comprises bundle of His, the left and right bundle branches, and Purkinje fibers; this subset of cardiomyocytes are distinct from the working myocardium that supports contraction [10]. Gap junctions between cardiomyocytes contribute to rapid conduction of action potentials both within the VCS and in working myocardium. Purkinje fibers relay the action potential from the proximal VCS to the working myocardium and coordinate simultaneous contraction of the ventricular working myocardium. Connexons, made up of connexin protein subunits, create the pores between cardiomyocytes at gap junctions [11]. Cardiomyocytes within the VCS highly express Connexin 40 (Cx40), while both VCS and working myocardium express Connexin 43 (Cx43) [12]. Abnormal expression of either Cx40 or Cx43 have been reported in patients with ventricular conduction disease, and mouse models support roles for both connexins in normal ventricular conduction [13-16]. Of note, Purkinje fibers originate from trabecular myocardium, and how the persistence of trabecular myocardium affects the VCS and working myocardium is not well understood. In humans, left ventricular noncompaction cardiomyopathy (LVNC) is often associated with ventricular arrhythmias, but mechanisms remain unclear [17, 18]. Finally, structural abnormalities of the working myocardium, such as cardiac fibrosis, can cause arrhythmias due to disrupted conduction pathways and late depolarization within regions of fibrosis [19, 20].

Here, we describe the phenotype of adult mice that survived cardiomyocyte-specific excision of *S1pr1* during development. All mutant mice displayed cardiac hypertrabeculation consistent with ventricular noncompaction, which established this genetic approach as a model system for human left ventricular noncompaction. Mutant mice also showed prolonged QRS duration, a sign of aberrant ventricular conduction. Because the VCS is derived from trabecular myocardium, we asked whether the conduction system develops normally in hearts with abnormally persistent trabeculation.

## METHODS

### Nomenclature

Sphingosine-1-phosphate receptor 1 protein is called S1P_1_ by International Union of Basic and Clinical Pharmacology (IUPHAR) nomenclature. *S1pr1* is the mouse gene that encodes S1P_1_ [1].

### Mouse strains

Richard Proia (National Institute of Diabetes and Digestive and Kidney Diseases, Bethesda, USA) provided *S1pr1* global heterozygous knockout mice (*S1pr1*^*tm2Rlp*^, designated as *S1pr1*^+/-^ mice) [3]. Dr. Jerold Chun provided mice carrying a floxed *S1pr1* allele (*S1pr1*^*tm1Jch*^; designated as *S1pr1*^f/f^) [5]. Dr. Xu Peng (Scott & White Memorial Hospital, Temple, USA) provided *Mlc2a*-Cre knockin mice (*Myl7*^*tm1(cre)Krc*^, designated as *Mlc2a*^Cre/+^) [21]. Mice with conditional deletion of *S1pr1* were generated as previously described [6]. We used *Mlc2a*^Cre/+^ rather than *Mlc2v*^Cre/+^ due to more efficient Cre-mediated excision in embryonic ventricular cardiomyocytes using *Mlc2a*^Cre/+^ [6, 22]. Briefly, to avoid germline excision in the setting of exogenous Cre expression during development, we maintained *Mlc2a*^Cre/+^ on the *S1pr1*^+/-^ background. Conditional *S1pr1* mutant mice were generated in crosses of *S1pr1*^f/f^ females (mixed C57BL6/J and 129SvJ strain background) with *Mlc2a*^Cre/+^;*S1pr1*^+/-^ males (C57BL6/J strain). This breeding strategy created *Mlc2a*^Cre/+^;*S1pr1*^f/-^ mutant mice that lack *S1pr1* in cardiomyocytes, as well as *S1pr1*^f/+^, *S1pr1*^f/-^, and *Mlc2a*^Cre/+^;*S1pr1*^f/+^ littermate controls (Figure 1A). Both male and female mice were used between 6-12 months of age in adult mouse echocardiographic and electrocardiographic studies. Dr. Shaun Coughlin (University of California, San Francisco) provided sphingosine kinase 1 global knockout mice (*Sphk1*^tm1Cgh^, designated as *Sphk1*^-/-^), conditional sphingosine kinase 1 mice (*Sphk1*^tm2Cgh^, designated as *Sphk1*^f/f^), and sphingosine kinase 2 global knockout mice (*Sphk2*^tm1.1Cgh^, designated as *Sphk2*^-/-^). For *Sphk1;Sphk2* cardiomyocyte deletion studies, we maintained *Mlc2a*^Cre/+^ on the *Sphk1*^+/-^; *Sphk2*^-/-^ background. *Mlc2a*^Cre/+^; *Sphk1*^+/-^; *Sphk2*^-/-^ males were bred with *Sphk1*^f/f^; *Sphk2*^-/-^ females to generate *Mlc2a*^Cre/+^; *Sphk1*^f/-^; *Sphk2*^-/-^ with deletion of sphingosine kinases in cardiomyocytes (Supplemental Figure 1A).

**Figure 1.**
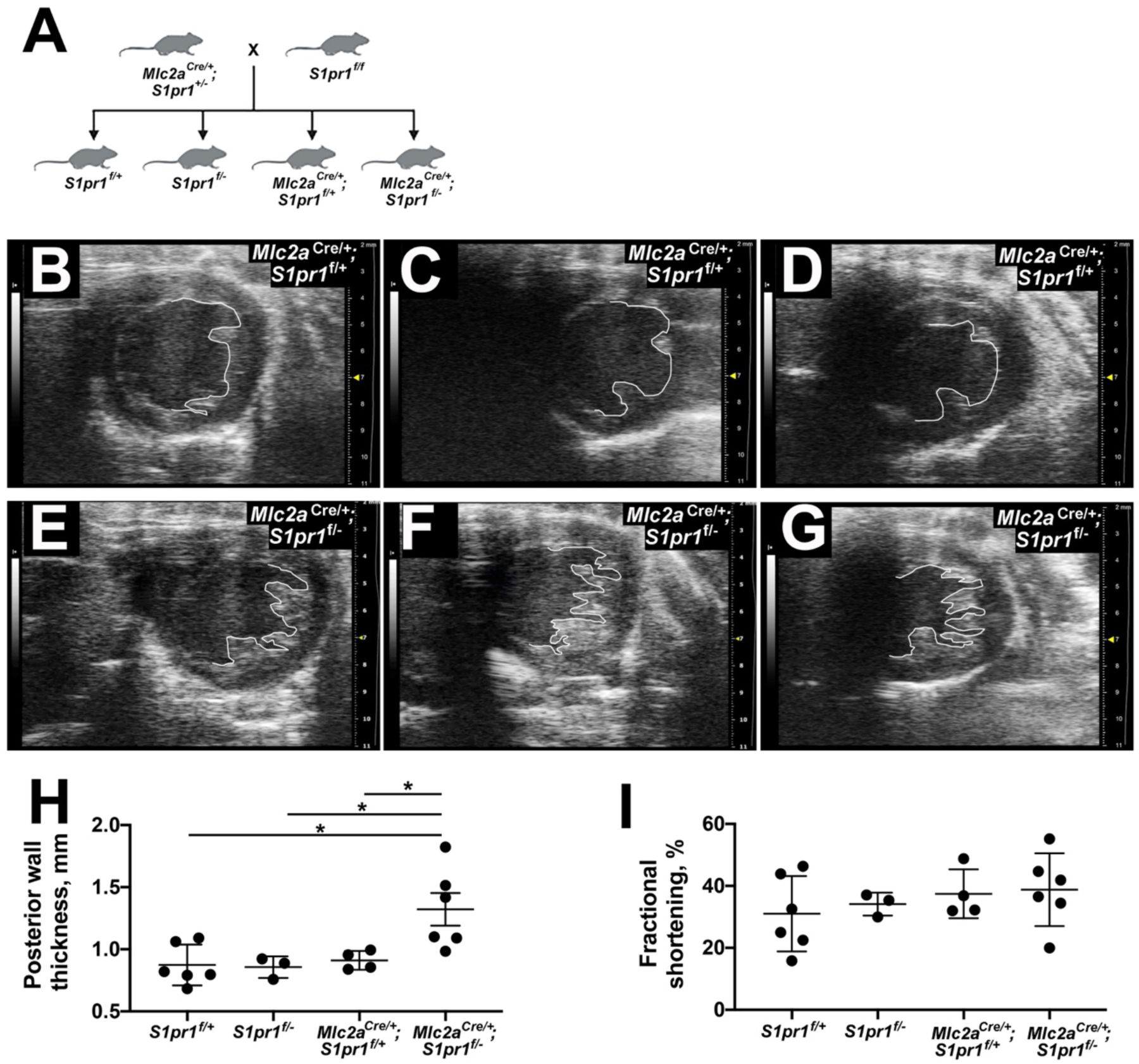
Cardiomyocyte-specific excision of *S1pr1* during development leads to abnormal cardiac structure in adult mice. A) Breeding strategy. (B-G) Representative short-axis echocardiography images from *Mlc2a*^*Cre/+*^;*S1pr1*^f/+^ control (B-D) and *Mlc2a*^*Cre/+*^;*S1pr1*^f/-^ mutant (E-G) mice. The white line highlights the endocardial border. Note hypertrabeculation in mutant hearts. (H) Posterior wall thickness in mice from the four genotypes generated by the breeding strategy. *, p < 0.05 by one-way ANOVA with Bonferroni post-hoc test for multiple comparisons. (I) Fractional shortening in mice from the four genotypes revealed no significant differences among the four genotypes. n = 3-6 mice per genotype.

### Mouse procedures

All mouse procedures were approved by the Institutional Animal Care and Use Committee at Northwestern University.

#### Embryo collection

The morning of vaginal plug was defined as 0.5 days post coitus (dpc). Pregnant females were euthanized at 15.5 dpc. Embryos were removed to ice-cold PBS, dissected away from the placenta, and snap frozen in OCT using liquid nitrogen-cooled isopentane. For a subset of embryos, the heart was dissected out and frozen in OCT. In all cases, a portion of the embryo was removed for genotyping.

#### Echocardiography

Adult mouse echocardiograms were recorded using a Vevo2100 echocardiography platform with a 550 MHz solid-state probe (FujiFilm, Toronto, ON, Canada). Mice were anesthetized using 2% isoflurane, and anesthesia was maintained using 1% isoflurane during collection of echocardiograms. Mouse heart rate was maintained between 400 and 600 beats per minute (bpm). Recordings were taken from both parasternal long-axis and short-axis views. Left ventricular wall thickness and internal dimensions during both systole and diastole were measured using a short-axis view approximately midway through the long-axis of the chamber. Fractional shortening was calculated using measurement from 3 beats per recording.

#### Electrocardiography

Adult mouse electrocardiograms (ECGs) were recorded using the ECGenie platform (Mouse Specifics, Inc., Framingham, MA). Mice were acclimated to the apparatus, then placed on the electrode pad for approximately 20 minutes. Mice were not anesthetized during the recording of ECGs. A consistent and representative subsection of 10-20 cardiac cycles of the ECG recordings was analyzed using the EzCG Signal Analysis Software.

#### Adult Heart Collection

Mice were anesthetized using 4% isoflurane. Once anesthetized, thoracotomy was performed, heart was fully exposed, the inferior vena cava was severed to release blood, and 3-5 milliliters of cardioplegia (128 mM sodium chloride, 2.5 mM magnesium sulfate, 15 mM potassium chloride, 10 mM glucose, 0.62 mM sodium dihydrate phosphate, 10mM HEPES, 1mM calcium chloride dihydrate) solution was perfused into the left ventricle (LV). The heart was removed, placed into an aluminum ladle, and snap frozen using liquid nitrogen. Hearts were stored at −80 C until sectioning.

#### Cryosectioning and Immunofluorescence staining

Cryosections of 10μm thickness were collected as frontal sections of the hearts. Sections selected for staining contained both ventricles and the mitral valve in order to include the ventricular conduction system (VCS). Immunostaining was performed as previously described [6, 23]. Briefly, sections were fixed using acetone and permeabilized using Triton X-100. Sections were blocked in Western Blocking Reagent (Roche #11921673001) and a second blocking step with goat anti-mouse IgG (H+L) monovalent Fab fragment (Jackson ImmunoResearch #115-007-003) was used to reduce background in the setting of mouse monoclonal primary antibodies. Sections were incubated with primary antibody overnight at 4 degrees Celsius. Primary antibodies included anti-α-actinin (sarcomeric) (1:2000, Sigma #A7811), anti-Connexin-40 (1:1500, Alpha Diagnostic #Cx40-A), anti-Connexin-43 (1:1500, Sigma #C6219), and anti-N-Cadherin (1:300, Abcam #ab18203). Secondary antibodies included goat anti-mouse Alexa Fluor 488 (1:2000, Life Technologies #A-11001) and goat anti-rabbit Alexa Fluor 568 (1:1000, Life Technologies #A-11011). Nuclei were stained using Hoechst (1:2000, Life Technologies #H3570). Slides were post-fixed using 1% PFA and mounted using Prolong Diamond Anti-Fade (ThermoFisher #P36961).

#### Microscopy

Imaging was performed using a Zeiss Axio Observer epifluorescence microscope and ZEN imaging software, and 40X images were obtained with an Apotome-2. Within each experiment, images were taken using the same exposure times per channel for each sample. Composite images of entire sections were taken at 10X magnification using the tiling function within ZEN. Images were processed using Fiji [24]; within each experiment, brightness, contrast, and thresholding were performed at the same values for each image.

#### Immunoblotting

Ventricular tissue was collected at seven weeks of age and homogenized in lysis buffer (150 mM sodium chloride, 1.0% Triton X-100, 0.5% sodium deoxycholate, 0.1% Sodium dodecyl sulfate, 50 mM Tris, pH 8.0) using silica beads and a homogenizer (BioSpec). Lysates were measured using the Pierce BCA assay kit (Thermo Scientific #23225) and normalized to 10 μg total protein. Lysates were separated using polyacrylamide gel electrophoresis and gel was transferred to nitrocellulose (NC) membranes. Membranes were blocked in 5% BSA prepared in 1x TBS-T. After blocking, NC membranes were incubated with primary antibodies Connexin 40 (1:1000, Cx40-A, Alpha Diagnostic Intl Inc.), Connexin 43 (1:1000, Cx43, Sigma C6219), or Contactin 2 (1:1000, R&D Systems, AF4439). Membranes were incubated with secondary antibody Goat anti-rabbit IgG-HRP (1:1000, Santa Cruz Biotechnology sc-2004) or Donkey anti-goat IgG-HRP (1:1000, Santa Cruz Biotechnology sc-2020). Signals were detected using SuperSignal West Pico PLUS Chemiluminescent Substrate (Thermo Scientific #34577). Total protein was determined using Memcode Reversible Protein Stain Kit (Thermo Scientific #24580).

#### Statistics

All measurements were done blind to genotype and treatment. Differences were assessed using one-way ANOVA, and post-hoc individual comparisons were performed using the Bonferroni test. All statistical analyses were performed using GraphPad software (La Jolla, CA). P < 0.05 was considered statistically significant.

## RESULTS

### Cardiomyocyte-specific excision of *S1pr1* during development leads to abnormal cardiac structure in adult mice

We generated mice with cardiomyocyte-specific deletion of *S1pr1* during embryonic development using a Cre recombinase driven by the promoter for *Myl7* (encodes myosin light chain 2a or Mlc2a, hereafter called *Mlc2a*^Cre/+^). To avoid germline excision in the setting of exogenous Cre expression during development, we maintained *Mlc2a*^Cre/+^ on the *S1pr1*^+/-^ background. Conditional *S1pr1* mutant mice were generated in crosses of *S1pr1*^f/f^ females with *Mlc2a*^Cre/+^;*S1pr1*^+/-^ males. This breeding strategy created *Mlc2a*^Cre/+^;*S1pr1*^f/-^ mutant mice that lack *S1pr1* in cardiomyocytes, as well as *S1pr1*^f/+^, *S1pr1*^f/-^, and *Mlc2a*^Cre/+^;*S1pr1*^f/+^ littermate controls (Figure 1A). We previously reported that *Mlc2a*^Cre/+^; *S1pr1*^f/-^ mice show cardiac structural abnormalities including ventricular noncompaction [6]. The majority (68%) of mutant mice died in the perinatal period, but the surviving cardiomyocyte *S1pr1* mutant animals had a normal lifespan of 18 months or longer. Echocardiography revealed that hearts from control littermates showed normal left ventricular wall thickness (Figure 1B-D and Supplemental Movies 1-3); by contrast, hearts from *Mlc2a*^Cre/+^; *S1pr1*^f/-^ mutant mice displayed persistent trabeculation consistent with ventricular noncompaction (Figure 1E-G and Supplemental Movies 4-6). Left ventricular posterior wall thickness was also increased in mutant hearts; on visual inspection, increased posterior wall thickness corresponded to increased ventricular trabeculation (Figure 1H). Unexpectedly, left ventricular systolic function remained normal in cardiomyocyte *S1pr1* mutant mice (Figure 1I). None of the adult *Mlc2a*^Cre/+^; *S1pr1*^f/-^ mutant mice had a ventricular septal defect, but all demonstrated abnormal trabeculation.

The source of S1P ligand that activates cardiomyocyte S1P_1_ is not definitively known and potentially includes endocrine, paracrine, and autocrine sources. Mammals carry two sphingosine kinase genes, sphingosine kinase-1 (*Sphk1*) and sphingosine kinase-2 (*Sphk2*); their protein products catalyze phosphorylation of sphingosine to S1P [25, 26]. To assess whether S1P produced within cardiomyocytes is necessary for the hypertrabeculation phenotype, we generated mice with cardiomyocyte-specific deletion of *Sphk1* and *Sphk2*. For these studies, we maintained *Mlc2a*^Cre/+^ on the *Sphk1*^+/-^; *Sphk2*^-/-^ background. *Mlc2a*^Cre/+^; *Sphk1*^+/-^; *Sphk2*^-/-^ males were bred with *Sphk1*^f/f^; *Sphk2*^-/-^ females to generate *Mlc2a*^Cre/+^; *Sphk1*^f/-^; *Sphk2*^-/-^ mice with deletion of sphingosine kinases in cardiomyocytes (Supplemental Figure 1A). At weaning, we found the expected distribution of all genotypes including *Mlc2a*^Cre/+^; *Sphk1*^f/-^; *Sphk2*^-/-^ (Supplemental Table 1). Furthermore, we did not observe differences in cardiac structure by echocardiography in *Mlc2a*^Cre/+^; *Sphk1*^f/-^; *Sphk2*^-/-^ versus littermate control mice (Supplemental Figure 1B and 1C). These results argue against autocrine or cardiomyocyte cell-autonomous roles for S1P in cardiac development; rather, these results indicate that either endocrine or paracrine sources of S1P act on the S1P_1_ receptor in cardiomyocytes.

### Cardiomyocyte *S1pr1* mutants display abnormal ventricular conduction

During echocardiography, we frequently noted wide QRS intervals on the electrocardiography (ECG) tracings in mice that also showed features of hypertrabeculation on echocardiography imaging (Figure 2A). The QRS interval reflects ventricular depolarization, and multiple forms of cardiomyopathy are associated with prolonged QRS interval [27]. To further investigate whether cardiomyocyte-specific *S1pr1* mutant mice had prolonged QRS, and to determine whether this feature was dependent or independent of isoflurane anesthesia, we collected ECG data in awake mice using the ECGenie platform. We observed no difference in QRS duration between *S1pr1*^f/+^ and *S1pr1*^f/-^ mice, intermediate QRS prolongation in *Mlc2a*^*Cre/+;*^ *S1pr1*^*f/+*^ mice, and the longest QRS duration in *Mlc2a*^*Cre/+*^; *S1pr1*^*f/-*^ mutant mice (Figure 2B). QRS duration was significantly longer in *Mlc2a*^*Cre/+*^; *S1pr1*^*f/-*^ mutant mice as compared with *S1pr1*^f/+^ and *S1pr1*^f/-^ mice, but the intermediate QRS duration in *Mlc2a*^*Cre/+*^; *S1pr1*^*f/+*^ control mice was not significantly different from any other group. These results suggest a potential interaction between *Mlc2a* haploinsufficiency and *S1pr1* dosage. The lack of difference in QRS duration between *S1pr1*^f/+^ mice and *S1pr1*^f/-^ mice indicates that heterozygosity for *S1pr1* alone does not drive changes in cardiac conduction. However, these data suggest that the loss one copy of *Mlc2a*, the presence of the Cre recombinase, or both may contribute to ventricular conduction abnormalities, and that the combination of these genetic changes in addition to loss of *S1pr1* in cardiomyocytes further compounds ventricular conduction abnormalities.

**Figure 2.**
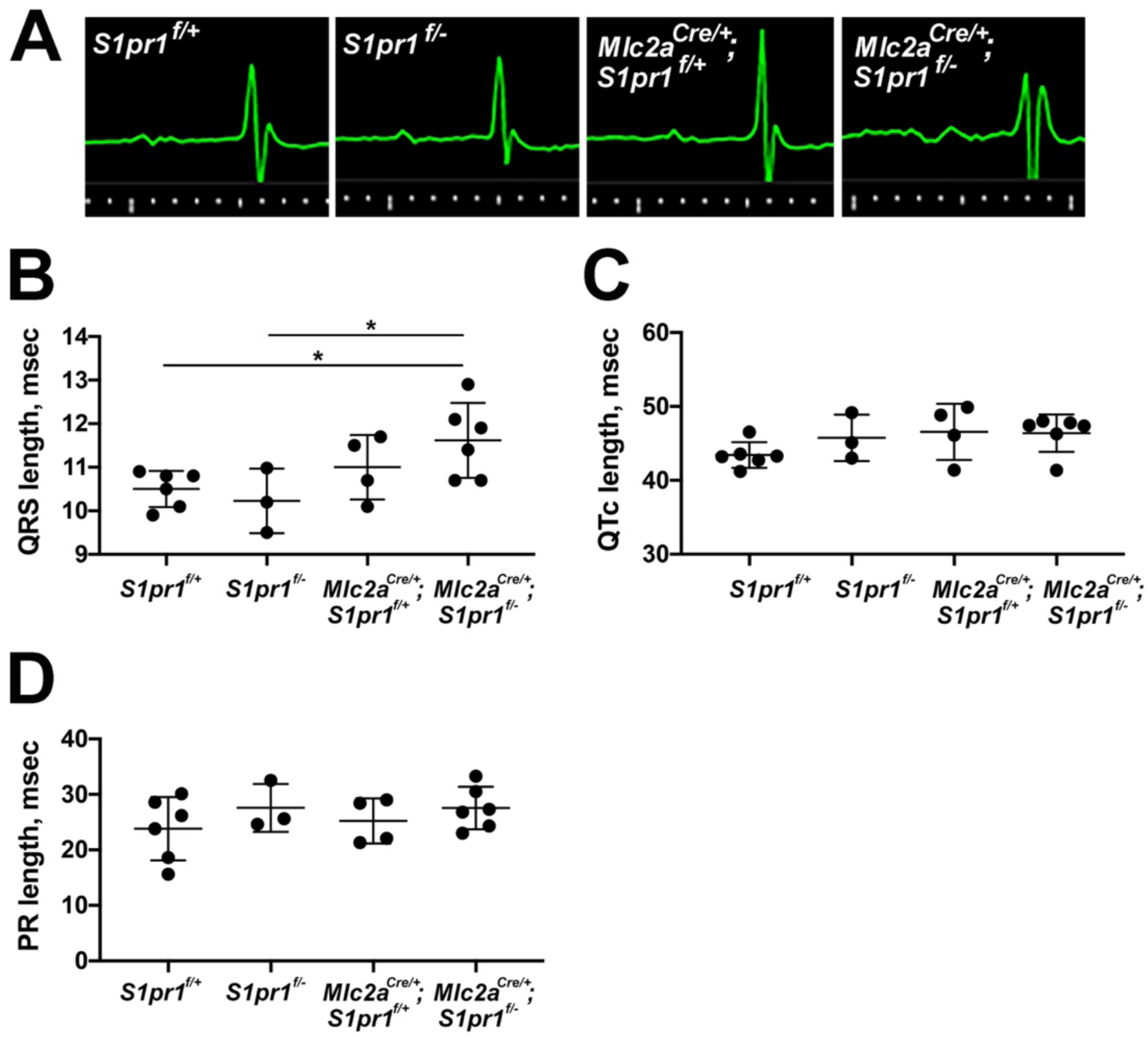
Cardiomyocyte *S1pr1* mutants display abnormal ventricular conduction. A) Representative electrocardiography (ECG) tracings in green obtained during echocardiography. Genotypes are noted for each tracing. B) QRS length, C) Corrected QT (QTc) length, and D) PR length from mice of each genotype as measured in awake mice using the ECGenie system. n = 3-6 mice per genotype. *, p < 0.05 by one-way ANOVA with Bonferroni post-hoc test for multiple comparisons.

Given the genotype-specific differences in ventricular depolarization, we looked for changes in other portions cardiac conduction pathway. We observed no differences in the corrected QT (QTc) interval, which reflects ventricular repolarization (Figure 2C). We also found no genotype-specific differences in the PR interval, which reflects conduction from the atria to the ventricles through the atrioventricular node (Figure 2D). Overall, these results indicate that cardiomyocyte-specific *S1pr1* deletion specifically affects ventricular depolarization during the cardiac conduction cycle.

### Ventricular conduction system development after embryonic cardiomyocyte *S1pr1* deletion

During heart development, Purkinje fibers originate from trabecular myocardium. Like other components of the VCS, Purkinje fibers highly express Connexin 40 (Cx40, encoded by *Gja5*). Furthermore, altered expression of Cx40 and Connexin 43 (Cx43) have been observed in mouse and human cardiomyopathies with aberrant ventricular conduction [13-16]. To investigate this potential mechanism of abnormal ventricular conduction in cardiomyocyte *S1pr1* mutant mice, we first assessed Cx40 expression in hearts from mice at 15.5 dpc. As previously reported [6], s-α-actinin staining of embryonic hearts from the three control genotypes showed normal ventricular compaction (Figure 3A-C), with roughly equal thickness of the trabecular and compact myocardium; by contrast, embryonic hearts from *Mlc2a*^*Cre/+*^; *S1pr1*^*f/-*^ mutant mice displayed deep trabeculations and a thin compact wall (Figure 3D). As expected, Cx40 was expressed in atria and trabecular myocardium in the three control genotypes (Figure 3A’-C’). In *Mlc2a*^*Cre/+*^; *S1pr1*^*f/-*^ mutant hearts, we predicted that all trabecular myocardium would express Cx40 at a high level. However, Cx40 expression was not uniformly expressed in the trabecular myocardium of *Mlc2a*^*Cre/+*^; *S1pr1*^*f/-*^ mutant hearts; trabecular myocardium closer to the thin compact wall expressed lower levels of Cx40 by immunostaining (Figure 3D’). This Cx40 expression pattern has been noted in a subset of ventricular noncompaction models and has been termed “intermediate myocardium” [28, 29]. We observed no significant differences in the immunostaining patterns for Connexin 43 (Cx43) or N-Cadherin (Supplemental Figure 2).

**Figure 3.**
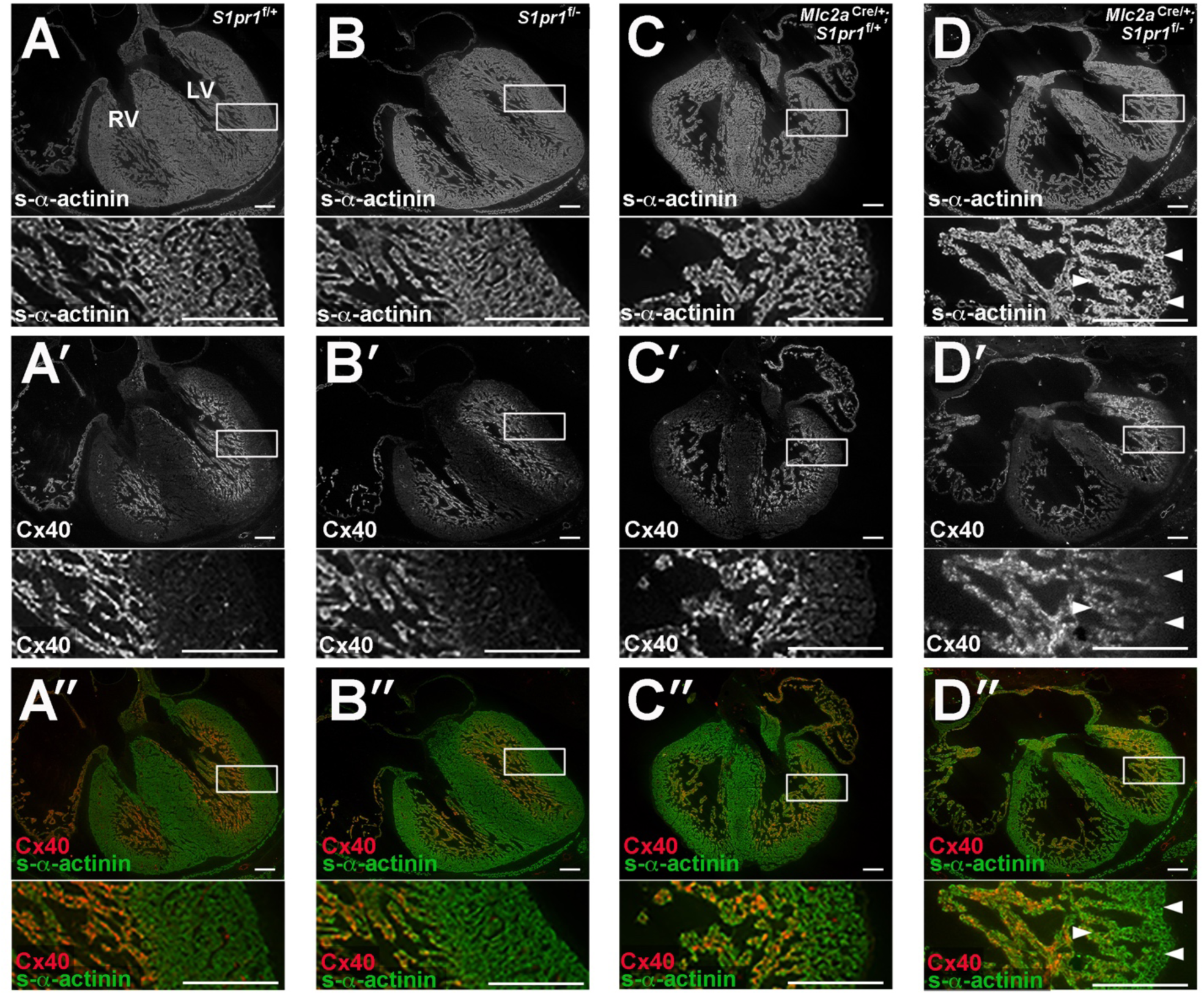
Connexin 40 (Cx40) expression pattern is disrupted in mice with embryonic cardiomyocyte *S1pr1* deletion. Embryos were collected at 15.5 dpc. A) *S1pr1*^f/+^ embryonic heart. B) *S1pr1*^f/-^ embryonic heart. C) *Mlc2a*^*Cre/+;*^ *S1pr1*^*f/+*^ embryonic heart. D) *Mlc2a*^*Cre/+;*^ *S1pr1*^*f/-*^ mutant embryonic heart. A-D) Immunostaining for s-α-actinin to mark all cardiomyocytes. A’-D’) Immunostaining for Cx40. A’’-D’’) Merged images for s-α-actinin and Cx40. Boxes indicate magnified region shown below each low-power image. Arrows highlight regions of reduced Cx40 expression in trabecular myocardium adjacent to the thin compact layer in the mutant embryonic heart. Representative images from n = 3 per genotype are shown. LV, left ventricle. RV, right ventricle. Scale bar, 200 μm.

Next, we examined Cx40 expression patterns in adult mice. Hearts from *Mlc2a*^*Cre/+*^; *S1pr1*^*f/-*^ mutant mice showed abnormal structure (Figure 4; of note, the heart shown in Figure 4B is the same as in Figure 1F and Supplemental Movie 5). Unlike the abnormal Cx40 expression pattern in embryonic *Mlc2a*^*Cre/+*^; *S1pr1*^*f/-*^ mutant hearts, adult mutant hearts demonstrated a Cx40 expression pattern that was similar to controls and restricted to cardiomyocytes within the bundle branches and Purkinje fibers. We did not observe an increase of Cx40 immunostaining in hypertrabeculated myocardium; rather, high-power imaging suggested Cx40 expression in fewer cardiomyocytes along the left bundle branch in *Mlc2a*^*Cre/+*^; *S1pr1*^*f/-*^ mutant hearts (Figure 4B). To quantitatively assess Cx40 expression, we collected whole hearts and isolated protein from combined left and right ventricles for immunoblotting. We did not detect a significant difference in Cx40 levels in ventricular lysates between *S1pr1*^f/+^ and *Mlc2a*^*Cre/+*^; *S1pr1*^*f/-*^ mutant mice (Figure 4C). Because Cx40 is also expressed in coronary artery endothelium, we further assessed expression of Contactin-2 (Cntn2), a cell adhesion molecule expressed specifically in postnatal VCS cardiomyocytes [30]. Again, we observed no differences in ventricular Cntn2 expression levels between control and mutant mice (Figure 4D). Finally, we assessed Cx43 and NCad expression via immunofluorescence, and similar patterns and intensities were noted across genotypes (Supplemental Figure 3). Together, these results suggest that the noncompaction and abnormal Cx40 expression pattern in *Mlc2a*^*Cre/+;*^ *S1pr1*^*f/-*^ mutant embryos may not disrupt development and maintenance of the VCS at later stages.

**Figure 4.**
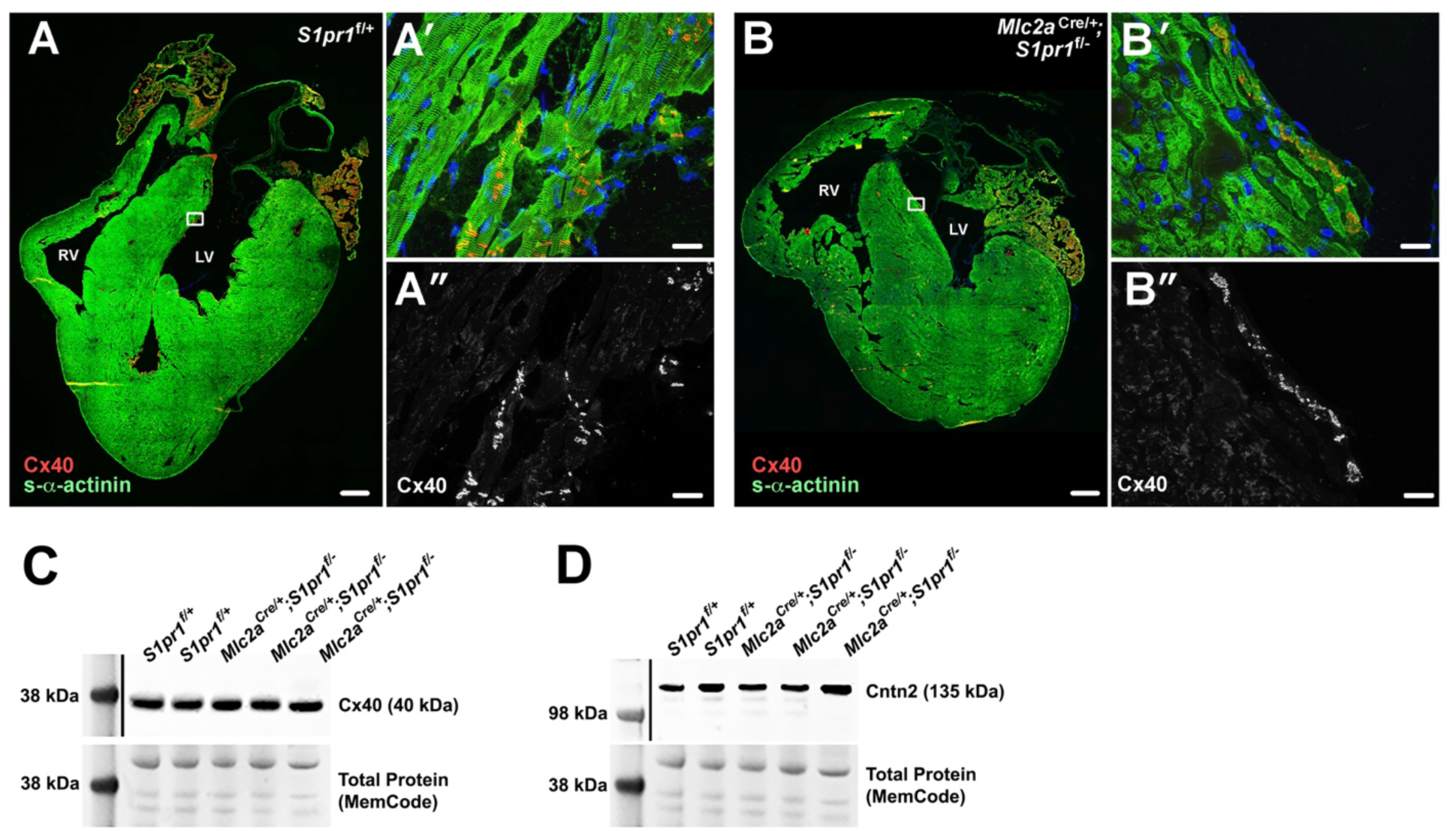
Normal markers of the ventricular conduction system in mice with embryonic cardiomyocyte *S1pr1* deletion. Adult hearts were perfused with cardioplegia solution, snap frozen, cryosectioned, and immunostained for s-α-actinin and Cx40. Representative images from n = 3-6 mice per genotype are shown. A) *S1pr1*^f/+^ control heart tiled imaged to show entire section. Scale bar, 500 μm. A’) High-power image from boxed area in panel A. Note Cx40 staining at intercalated disks of VCS cardiomyocytes. Scale bar, 20 μm. A’’) Cx 40 staining alone from the high-power image. B) *Mlc2a*^*Cre/+;*^ *S1pr1*^*f/-*^ mutant heart tiled image. This heart is also represented in Figure 1F and Supplemental Movie 5. Scale bar, 500 μm. B’) High-power image from boxed area in panel B. Note Cx40 staining in fewer VCS cardiomyocytes as compared to B’. Scale bar, 20 μm. B’’) Cx40 staining alone from the high-power image. C) Immunoblot of lysates from whole left and right ventricles show no difference in Cx40 levels between *S1pr1*^f/+^ control and *Mlc2a*^*Cre/+;*^ *S1pr1*^*f/-*^ mutant hearts. D) Immunoblot of lysates from whole left and right ventricles show no difference in Contactin 2 (Cntn2) levels between *S1pr1*^f/+^ control and *Mlc2a*^*Cre/+;*^ *S1pr1*^*f/-*^ mutant hearts.

### Cardiomyocyte *S1pr1* mutants display abnormal increased cardiac fibrosis

Structural abnormalities of the working myocardium, such as cardiac fibrosis, can also disrupt ventricular conduction. To investigate cardiac fibrosis as a potential mechanism for QRS prolongation in cardiomyocyte *S1pr1* mutants, we immunostained heart sections for Collagen 1a1 (Col1a1). We detected low levels of Col1a1 in heart sections from *S1pr1*^f/+^ and *S1pr1*^f/-^ mice, while sections from *Mlc2a*^*Cre/+*^; *S1pr1*^*f/+*^ mutant showed moderate levels of Col1a1 immunostaining (Figure 5A-C). Heart sections from *Mlc2a*^*Cre/+*^; *S1pr1*^*f/-*^ mutant mice displayed the highest level of Col1a1 signal (Figure 5D). We used thresholding to measure Col1a1 signal in multiple regions across heart sections and found significantly higher signal in *Mlc2a*^*Cre/+*^; *S1pr1*^*f/-*^ mutant hearts as compared with *S1pr1*^f/+^ and *S1pr1*^f/-^ mice (Figure 5E). As we observed with QRS duration, the difference in Col1a1 signal did not reach statistical significance between *Mlc2a*^*Cre/+*^; *S1pr1*^*f/+*^ hearts and *Mlc2a*^*Cre/+*^; *S1pr1*^*f/-*^ mutant hearts. These results demonstrate that cardiomyocyte-specific loss of *S1pr1* increases cardiac fibrosis in surviving adult mice, and that this increased cardiac fibrosis is the most likely cause of abnormal ventricular conduction in these mutant mice.

**Figure 5.**
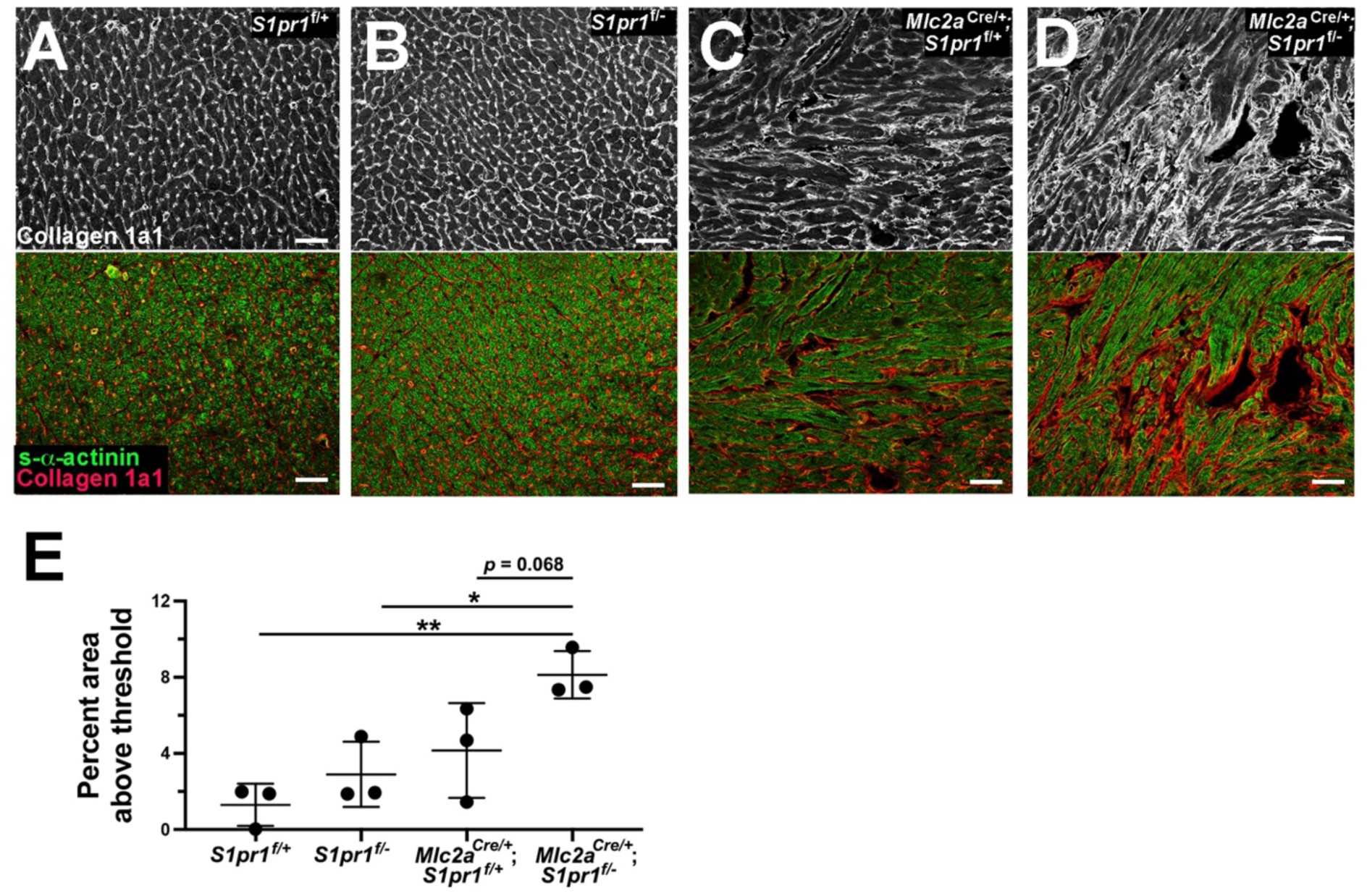
Cardiomyocyte *S1pr1* mutants display increased Collagen 1a1 fibrosis marker. A-D) Adult hearts were perfused with cardioplegia solution, snap frozen, cryosectioned, and immunostained for Collagen 1a1 (Col1a1, top row) and s-α-actinin (merged image with Col1a1, bottom row). A) *S1pr1*^f/+^ heart. B) *S1pr1*^f/-^ heart. C) *Mlc2a*^*Cre/+;*^ *S1pr1*^*f/+*^ heart. D) *Mlc2a*^*Cre/+;*^ *S1pr1*^*f/-*^ mutant heart. E) Col1a1 signal was measured using the percent of the image area above threshold. 6 images across the left ventricle were taken for each heart blind to genotype, and percent area of Col1a1 signal above threshold was averaged. n = 3 per genotype. *, p < 0.05 and **, p < 0.01 by one-way ANOVA with Bonferroni post-hoc test for multiple comparisons.

## DISCUSSION

Here we report the phenotype of adult mice that survive embryonic deletion of *S1pr1* in cardiomyocytes. We observed cardiac hypertrabeculation consistent with ventricular noncompaction, preserved systolic function, and normal lifespan in surviving *Mlc2a*^*Cre/+*^; *S1pr1*^*f/-*^ mutant mice. Cardiomyocyte *S1pr1* mutant mice also demonstrated conduction abnormalities with prolonged QRS in the awake and active state. The conduction system marker Cx40 displayed an abnormal expression pattern in *Mlc2a*^*Cre/+*^; *S1pr1*^*f/-*^ mutant embryos. However, Cx40 and Cntn2 expression in adult mice did not differ from littermate controls, which suggests normalized development and maintenance of the VCS in cardiomyocyte *S1pr1* mutant mice. Finally, cardiomyocyte *S1pr1* mutant mice demonstrated increased cardiac fibrosis.

Purkinje cells are thought to arise from trabecular cardiomyocytes [31], and conduction system development in the setting of persistent trabeculation and noncompaction is not well understood. In mammals, the cardiac conduction system (CCS) is specified early in cardiac development. The conduction system marker CCS-LacZ revealed that CCS-specific expression begins in the sinoatrial node at 9.5 dpc, extends from atria to the early atrioventricular node and ventricular trabecular myocardium between 10.5 dpc and 12.5 dpc, and has the appearance of a full CCS with Purkinje fibers by 13.5 dpc [32]. Subsequent fate mapping studies in Cx40-positive cardiomyocytes showed that VCS progenitor cells differentiate into both mature conductive cells and working myocardial cells prior to 14.5 dpc, but that by 16.5 dpc the conductive fate is established in Cx40-positive VCS progenitor cells [10]. Experiments in *Cx40-* eGFP reporter mice demonstrated that *Nkx2*.*5* haploinsufficiency disrupted postnatal maturation of the Purkinje fiber network; furthermore, embryonic deletion of *Nkx2*.*5* in trabecular myocardium using *Cx40-CreERT2* led to hypertrabeculation (but normal compact wall thickness), Purkinje fiber hypoplasia, subendocardial fibrosis, and conduction abnormalities in adult hearts [33, 34]. Of note, the Cx40-eGFP expression pattern was normal in hearts from *Cx40-eGFP;Nkx2*.*5*^*+/-*^ embryos at 16.5 dpc, while Purkinje fiber morphology was disrupted at birth in *Cx40-eGFP;Nkx2*.*5*^*+/-*^ hearts [33]. By contrast, our results in 15.5 dpc *Mlc2a*^*Cre/+*^; *S1pr1*^*f/-*^ mutant embryos revealed that trabecular myocardium did not uniformly express Cx40, with lower expression in the region adjacent to the thin compact myocardium (Figure 3D). However, we did not observe differences in Cx40 expression pattern or level in adult *Mlc2a*^*Cre/+*^; *S1pr1*^*f/-*^ mutant hearts. Therefore, while this region of reduced Cx40 trabecular myocardium may contribute to abnormal compaction, the altered Cx40 expression in that region does not appear to affect further maturation of the VCS in adult mice. Future studies with *Cx40-eGFP* or *Cntn2-EGFP* lines will more definitively assess bundle branch density in *Mlc2a*^*Cre/+*^; *S1pr1*^*f/-*^ mutant hearts.

Other mouse models of ventricular noncompaction have demonstrated non-uniform expression of Cx40 in the trabecular myocardium; specifically, disruption of some components of the Notch signaling pathway (combined deletion of cardiomyocyte *Jagged1 and Jagged2*, as well as endothelial overexpression of *Manic Fringe)* and endothelial-specific deletion of *Ino80* led to the same Cx40 expression pattern that we observed [29, 35]. This “intermediate myocardium” represents a margin between trabecular and compact myocardium, with cardiomyocytes that display trabecular morphology but express molecular markers of compact/non-trabecular myocardium. *S1pr1* was not reported to be differentially expressed in the other models that display intermediate myocardium, and whether S1P_1_ signaling pathways intersect with Notch signaling or Ino80-mediated chromatin remodeling in developing cardiomyocytes remains to be determined.

In adult mice, we observed an intermediate increase in QRS prolongation and cardiac fibrosis in *Mlc2a*^*Cre/+*^; *S1pr1*^*f/+*^ mice, as compared with Cre-negative controls and *Mlc2a*^*Cre/+*^; *S1pr1*^*f/-*^ mutant mice. These results suggest a potential interaction between *S1pr1* dosage and the presence of *Mlc2a*^*Cre/+*^. Heterozygosity for *S1pr1* did not lead to a change in either QRS duration or cardiac fibrosis, which indicates that *S1pr1* dosage alone was not the primary driver of these phenotypes. *Mlc2a* is expressed in both atrial and ventricular cardiomyocytes up to embryonic day 11 and then restricted to atrial cardiomyocytes by embryonic day 12 in mice [36]. *Mlc2a* heterozygous mice survive with no obvious cardiac phenotype [37], so it was predicted that haploinsufficiency would not affect cardiac development. However, *Mlc2a* null mice did not survive beyond 11.5 dpc due to severely reduced atrial contractility and failure to progress through normal ventricular trabeculation and compaction [37]. It is possible that haploinsufficiency for *Mlc2a* in the *Mlc2a*^*Cre/+*^ background could lead to subtle but functionally important reduction in atrial contractility; in such a setting, *Mlc2a* haploinsufficiency might create a sensitized background in which *S1pr1* gene dosage could contribute to the phenotype. Specifically, *Mlc2a* haploinsufficiency and *S1pr1* heterozygosity in cardiomyocytes may lead to the intermediate phenotype in *Mlc2a*^*Cre/+*^; *S1pr1*^*f/+*^ mice, while *Mlc2a* haploinsufficiency and loss of *S1pr1* in cardiomyocytes drives the full noncompaction, QRS prolongation, and cardiac fibrosis phenotypes. We cannot rule out the possibility that cardiomyocyte-specific deletion of *S1pr1* may also require *Mlc2a* haploinsufficiency for the noncompaction phenotype. Because *Mlc2a* is not expressed in ventricular cardiomyocytes after 12 dpc, it is unlikely that toxicity from accumulated Cre makes a large contribution to these phenotypes.

Excision of many different genes can lead to ventricular noncompaction; however, most of these genetic models are embryonic lethal with 100% penetrance [18]. Survival in a subset of cardiomyocyte *S1pr1* mutant mice indicates that this genetic approach can be used as a model for left ventricular noncompaction in humans. Humans with hypertrabeculation/left ventricular noncompaction are at risk for both systolic dysfunction and arrhythmia, and QRS prolongation in the setting of normal ejection fraction has been reported [17, 38]. While cardiomyocyte *S1pr1* mice do not develop systolic dysfunction, few adult mouse models exist for LVNC, and this will be a useful system to investigate this cardiomyopathy.

In terms of genetic cardiomyopathies, pathogenic *S1PR1* variants have not been described in humans. However, the Genome Aggregation Database (gnomAD), a data set of 125,748 exome sequences and 15,708 whole-genome sequences from unrelated individuals, identified two unique frameshift variant alleles at amino acid positions 255 and 256 (rs1166435525 and rs1395550411 in the dbSNP database, respectively) [39, 40]. These two variants disrupt S1P_1_ at the origin of transmembrane domain six. One stop-gained variant allele at amino acid position 283 was observed as well (rs760714388 in dbSNP); this nonsense variant terminates the S1P_1_ protein at extracellular domain four, between transmembrane domains six and seven. Furthermore, significantly fewer *S1PR1* missense variants were observed in gnomAD than expected (232 missense variants expected, 120 observed missense variants, z = 2.61). Finally, missense variants identified in the NHLBI Exome Sequencing Project have been functionally assessed: the Arg120Pro variant (rs149198314; identified in 1/13,005 alleles) failed to activate S1P_1_ signaling or internalization in response to S1P, and Arg120 was found to be critical for S1P binding [41-43]. Hence, individuals who are heterozygous for predicted loss-of-function *S1PR1* alleles have been observed at very low frequency, and cardiomyopathy exome/genome sequencing studies may reveal additional *S1PR1* variants that disrupt S1P1 function. Cardiac fibrosis has been associated with conduction abnormalities, including QRS prolongation, presumably due to the disruption of cardiomyocyte-cardiomyocyte electrical coupling and late depolarization in regions of fibrosis [19, 20]. Given the observation of both intermediate fibrosis and intermediate QRS prolongation in *Mlc2a*^*Cre/+*^; *S1pr1*^*f/+*^ mice that have no hypertrabeculation, we hypothesize that *Mlc2a* haploinsufficiency and *S1pr1* heterozygosity in cardiomyocytes can lower the threshold for cardiac fibrosis, possibly due to abnormal cardiac mechanics. The hypertrabeculation in *Mlc2a*^*Cre/+*^; *S1pr1*^*f/-*^ mutant mice may further impair cardiac mechanics and lead to increased fibrosis. This increase in fibrosis is the likely etiology of QRS prolongation in cardiomyocyte *S1pr1* mutant mice. In endothelial cells, S1P_1_ contributes to mechanosensing and endothelial cell alignment [9]; future experiments will investigate possible roles for S1P_1_ in cardiomyocyte mechanosensation pathways.

The source of S1P that activates cardiomyocyte S1P_1_ is not definitively known and potentially includes endocrine, paracrine, and autocrine sources. We found no phenotype in mice that lacked *Sphk1* and *Sphk2* in cardiomyocytes of the developing heart, which argues against autocrine S1P mechanism in cardiomyocytes and suggests that cardiomyocyte-specific generation of S1P is not necessary for normal cardiac development. Recent studies of Nogo-B, a protein that inhibits the rate-limiting enzyme in sphingolipid biosynthesis, suggest endothelial cells as a very likely origin [44].

In conclusion, we report that mice that survive embryonic deletion of *S1pr1* in cardiomyocytes show hypertrabeculation with normal lifespan and normal systolic function, prolonged QRS duration, and increased cardiac fibrosis. Our results suggest an interaction between *Mlc2a* haploinsufficiency (*Mlc2a*^Cre/+^) and *S1pr1* dosage in cardiomyocytes. Furthermore, few adult mouse models for hypertrabeculation/noncompaction exist, and *Mlc2a*^*Cre/+*^; *S1pr1*^*f/-*^ mutant mice represent a useful model to investigate this cardiomyopathy. The presence of conduction abnormalities in hypertrabeculated hearts, despite having normal systolic function, may be of clinical relevance for patients with hypertrabeculation.

## ACKNOWLEDGMENTS

This work was supported in part by NIH K08HL105657 and the Gilead Sciences Research Scholars Program in Cardiovascular Disease (LDW). The funding sources had no role in the design, collection and analysis of data, writing, or submission of the manuscript. The authors thank Dr. David Barefield, Dr. Elizabeth McNally, Dr. John Wasserstrom, Dr. Rishi Arora, and Dr. Gwendolyn Kaeser for editorial assistance.

## REFERENCES

[1] Y. Kihara, M. Maceyka, S. Spiegel, J. Chun, Lysophospholipid receptor nomenclature review: IUPHAR Review 8, Br J Pharmacol 171(15) (2014) 3575–94.

[2] J. Chun, Y. Kihara, D. Jonnalagadda, V.A. Blaho, Fingolimod: Lessons Learned and New Opportunities for Treating Multiple Sclerosis and Other Disorders, Annu Rev Pharmacol Toxicol 59 (2019) 149–170.

[3] Y. Liu, R. Wada, T. Yamashita, Y. Mi, C.X. Deng, J.P. Hobson, H.M. Rosenfeldt, V.E. Nava, S.S. Chae, M.J. Lee, C.H. Liu, T. Hla, S. Spiegel, R.L. Proia, Edg-1, the G protein-coupled receptor for sphingosine-1-phosphate, is essential for vascular maturation, J Clin Invest 106(8) (2000) 951–61.

[4] M. Matloubian, C.G. Lo, G. Cinamon, M.J. Lesneski, Y. Xu, V. Brinkmann, M.L. Allende, R.L. Proia, J.G. Cyster, Lymphocyte egress from thymus and peripheral lymphoid organs is dependent on S1P receptor 1, Nature 427(6972) (2004) 355–60.

[5] J.W. Choi, S.E. Gardell, D.R. Herr, R. Rivera, C.-W. Lee, K. Noguchi, S.T. Teo, Y.C. Yung, M. Lu, G. Kennedy, J. Chun, FTY720 (fingolimod) efficacy in an animal model of multiple sclerosis requires astrocyte sphingosine 1-phosphate receptor 1 (S1P1) modulation, Proc Natl Acad Sci USA 108(2) (2011) 751–756.

[6] H. Clay, L.D. Wilsbacher, S.J. Wilson, D.N. Duong, M. McDonald, I. Lam, K.E. Park, J. Chun, S.R. Coughlin, Sphingosine 1-phosphate receptor-1 in cardiomyocytes is required for normal cardiac development, Dev Biol 418(1) (2016) 157–65.

[7] M.L. Allende, T. Yamashita, R.L. Proia, G-protein-coupled receptor S1P1 acts within endothelial cells to regulate vascular maturation, Blood 102(10) (2003) 3665–7.

[8] K. Gaengel, C. Niaudet, K. Hagikura, B.L. Siemsen, L. Muhl, J.J. Hofmann, L. Ebarasi, S. Nyström, S. Rymo, L.L. Chen, M.-F. Pang, Y. Jin, E. Raschperger, P. Roswall, D. Schulte, R. Benedito, J. Larsson, M. Hellstrom, J. Fuxe, P. Uhlén, R. Adams, L. Jakobsson, A. Majumdar, D. Vestweber, A. Uv, C. Betsholtz, The Sphingosine-1-Phosphate Receptor S1PR1 Restricts Sprouting Angiogenesis by Regulating the Interplay between VE-Cadherin and VEGFR2, Dev. Cell 23(3) (2012) 587–599.

[9] B. Jung, H. Obinata, S. Galvani, K. Mendelson, B.-s. Ding, A. Skoura, B. Kinzel, V. Brinkmann, S. Rafii, T. Evans, T. Hla, Flow-Regulated Endothelial S1P Receptor-1 Signaling Sustains Vascular Development, Dev. Cell 23(3) (2012) 600–610.

[10] L. Miquerol, N. Moreno-Rascon, S. Beyer, L. Dupays, S.M. Meilhac, M.E. Buckingham, D. Franco, R.G. Kelly, Biphasic development of the mammalian ventricular conduction system, Circ Res 107(1) (2010) 153–61.

[11] S. Verheule, S. Kaese, Connexin diversity in the heart: insights from transgenic mouse models, Front Pharmacol 4 (2013) 81.

[12] N.J. Severs, A.F. Bruce, E. Dupont, S. Rothery, Remodelling of gap junctions and connexin expression in diseased myocardium, Cardiovasc Res 80(1) (2008) 9–19.

[13] T. Tang, N.C. Lai, A.T. Wright, M.H. Gao, P. Lee, T. Guo, R. Tang, A.D. McCulloch, H.K. Hammond, Adenylyl cyclase 6 deletion increases mortality during sustained beta-adrenergic receptor stimulation, J Mol Cell Cardiol 60 (2013) 60–7.

[14] D. Sedmera, J. Neckar, J. Benes, Jr., J. Pospisilova, J. Petrak, K. Sedlacek, V. Melenovsky, Changes in Myocardial Composition and Conduction Properties in Rat Heart Failure Model Induced by Chronic Volume Overload, Front Physiol 7 (2016) 367.

[15] E.E. Mueller, A. Momen, S. Masse, Y.Q. Zhou, J. Liu, P.H. Backx, R.M. Henkelman, K. Nanthakumar, D.J. Stewart, M. Husain, Electrical remodelling precedes heart failure in an endothelin-1-induced model of cardiomyopathy, Cardiovasc Res 89(3) (2011) 623–33.

[16] F.G. Akar, D.D. Spragg, R.S. Tunin, D.A. Kass, G.F. Tomaselli, Mechanisms underlying conduction slowing and arrhythmogenesis in nonischemic dilated cardiomyopathy, Circ Res 95(7) (2004) 717–25.

[17] E. Oechslin, R. Jenni, Left ventricular non-compaction revisited: a distinct phenotype with genetic heterogeneity?, Eur. Heart J. 32(12) (2011) 1446–1456.

[18] L. Wilsbacher, E.M. McNally, Genetics of Cardiac Developmental Disorders: Cardiomyocyte Proliferation and Growth and Relevance to Heart Failure, Annu Rev Pathol 11 (2016) 395–419.

[19] H.H. Nguyen, J.T. Wolfe, 3rd, D.R. Holmes, Jr., W.D. Edwards, Pathology of the cardiac conduction system in myotonic dystrophy: a study of 12 cases, J Am Coll Cardiol 11(3) (1988) 662–71.

[20] S. Nazarian, D.A. Bluemke, K.R. Wagner, M.M. Zviman, E. Turkbey, B.S. Caffo, M. Shehata, D. Edwards, B. Butcher, H. Calkins, R.D. Berger, H.R. Halperin, G.F. Tomaselli, QRS prolongation in myotonic muscular dystrophy and diffuse fibrosis on cardiac magnetic resonance, Magn Reson Med 64(1) (2010) 107–14.

[21] N. Wettschureck, H. Rütten, A. Zywietz, D. Gehring, T.M. Wilkie, J. Chen, K.R. Chien, S. Offermanns, Absence of pressure overload induced myocardial hypertrophy after conditional inactivation of Galphaq/Galpha11 in cardiomyocytes, Nat. Med. 7(11) (2001) 1236–1240.

[22] X. Peng, X. Wu, J.E. Druso, H. Wei, A.Y.-J. Park, M.S. Kraus, A. Alcaraz, J. Chen, S. Chien, R.A. Cerione, J.-L. Guan, Cardiac developmental defects and eccentric right ventricular hypertrophy in cardiomyocyte focal adhesion kinase (FAK) conditional knockout mice, Proc Natl Acad Sci USA 105(18) (2008) 6638–6643.

[23] L.D. Wilsbacher, S.R. Coughlin, Analysis of Cardiomyocyte Development using Immunofluorescence in Embryonic Mouse Heart, Journal of visualized experiments : JoVE (97) (2015).

[24] J. Schindelin, I. Arganda-Carreras, E. Frise, V. Kaynig, M. Longair, T. Pietzsch, S. Preibisch, C. Rueden, S. Saalfeld, B. Schmid, J.Y. Tinevez, D.J. White, V. Hartenstein, K. Eliceiri, P. Tomancak, A. Cardona, Fiji: an open-source platform for biological-image analysis, Nat Methods 9(7) (2012) 676–82.

[25] S. Spiegel, S. Milstien, Exogenous and intracellularly generated sphingosine 1-phosphate can regulate cellular processes by divergent pathways, Biochem Soc Trans 31(Pt 6) (2003) 1216–9.

[26] R. Pappu, S.R. Schwab, I. Cornelissen, J.P. Pereira, J.B. Regard, Y. Xu, E. Camerer, Y.W. Zheng, Y. Huang, J.G. Cyster, S.R. Coughlin, Promotion of lymphocyte egress into blood and lymph by distinct sources of sphingosine-1-phosphate, Science 316(5822) (2007) 295–8.

[27] A. Brenyo, W. Zareba, Prognostic significance of QRS duration and morphology, Cardiol J 18(1) (2011) 8–17.

[28] G. D’Amato, G. Luxan, G. Del Monte-Nieto, B. Martinez-Poveda, C. Torroja, W. Walter, M.S. Bochter, R. Benedito, S. Cole, F. Martinez, A.K. Hadjantonakis, A. Uemura, L.J. Jimenez-Borreguero, J.L. de la Pompa, Sequential Notch activation regulates ventricular chamber development, Nat Cell Biol 18(1) (2016) 7–20.

[29] S. Rhee, J.I. Chung, D.A. King, G. D’Amato, D.T. Paik, A. Duan, A. Chang, D. Nagelberg, B. Sharma, Y. Jeong, M. Diehn, J.C. Wu, A.J. Morrison, K. Red-Horse, Endothelial deletion of Ino80 disrupts coronary angiogenesis and causes congenital heart disease, Nat Commun 9(1) (2018) 368.

[30] B.A. Pallante, S. Giovannone, L. Fang-Yu, J. Zhang, N. Liu, G. Kang, W. Dun, P.A. Boyden, G.I. Fishman, Contactin-2 expression in the cardiac Purkinje fiber network, Circulation. Arrhythmia and electrophysiology 3(2) (2010) 186–94.

[31] S. Viragh, C.E. Challice, The development of the conduction system in the mouse embryo heart, Dev Biol 89(1) (1982) 25–40.

[32] S. Rentschler, D.M. Vaidya, H. Tamaddon, K. Degenhardt, D. Sassoon, G.E. Morley, J. Jalife, G.I. Fishman, Visualization and functional characterization of the developing murine cardiac conduction system, Development 128(10) (2001) 1785–92.

[33] S. Meysen, L. Marger, K.W. Hewett, T. Jarry-Guichard, I. Agarkova, J.P. Chauvin, J.C. Perriard, S. Izumo, R.G. Gourdie, M.E. Mangoni, J. Nargeot, D. Gros, L. Miquerol, Nkx2.5 cell-autonomous gene function is required for the postnatal formation of the peripheral ventricular conduction system, Dev Biol 303(2) (2007) 740–53.

[34] C. Choquet, T.H.M. Nguyen, P. Sicard, E. Buttigieg, T.T. Tran, F. Kober, I. Varlet, R. Sturny, M.W. Costa, R.P. Harvey, C. Nguyen, P. Rihet, S. Richard, M. Bernard, R.G. Kelly, N. Lalevee, L. Miquerol, Deletion of Nkx2-5 in trabecular myocardium reveals the developmental origins of pathological heterogeneity associated with ventricular non-compaction cardiomyopathy, PLoS Genet 14(7) (2018) e1007502.

[35] G. Luxán, J.C. Casanova, B. Martínez-Poveda, B. Prados, G. D’Amato, D. Macgrogan, A. Gonzalez-Rajal, D. Dobarro, C. Torroja, F. Martinez, J.L. Izquierdo-García, L. Fernández-Friera, M. Sabater-Molina, Y.-Y. Kong, G. Pizarro, B. Ibañez, C. Medrano, P. García-Pavía, J.R. Gimeno, L. Monserrat, L.J. Jiménez-Borreguero, J.L. de la Pompa, Mutations in the NOTCH pathway regulator MIB1 cause left ventricular noncompaction cardiomyopathy, Nat. Med. (2013).

[36] S.W. Kubalak, W.C. Miller-Hance, T.X. O’Brien, E. Dyson, K.R. Chien, Chamber specification of atrial myosin light chain-2 expression precedes septation during murine cardiogenesis, The Journal of biological chemistry 269(24) (1994) 16961–16970.

[37] C. Huang, F. Sheikh, M. Hollander, C. Cai, D. Becker, P.H. Chu, S. Evans, J. Chen, Embryonic atrial function is essential for mouse embryogenesis, cardiac morphogenesis and angiogenesis, Development 130(24) (2003) 6111–9.

[38] A. Frustaci, A. De Luca, V. Guida, T. Biagini, T. Mazza, C. Gaudio, C. Letizia, M.A. Russo, N. Galea, C. Chimenti, Novel alpha-Actin Gene Mutation p.(Ala21Val) Causing Familial Hypertrophic Cardiomyopathy, Myocardial Noncompaction, and Transmural Crypts. Clinical-Pathologic Correlation, Journal of the American Heart Association 7(4) (2018).

[39] M. Lek, K.J. Karczewski, E.V. Minikel, K.E. Samocha, E. Banks, T. Fennell, A.H. O’Donnell-Luria, J.S. Ware, A.J. Hill, B.B. Cummings, T. Tukiainen, D.P. Birnbaum, J.A. Kosmicki, L.E. Duncan, K. Estrada, F. Zhao, J. Zou, E. Pierce-Hoffman, J. Berghout, D.N. Cooper, N. Deflaux, M. DePristo, R. Do, J. Flannick, M. Fromer, L. Gauthier, J. Goldstein, N. Gupta, D. Howrigan, A. Kiezun, M.I. Kurki, A.L. Moonshine, P. Natarajan, L. Orozco, G.M. Peloso, R. Poplin, M.A. Rivas, V. Ruano-Rubio, S.A. Rose, D.M. Ruderfer, K. Shakir, P.D. Stenson, C. Stevens, B.P. Thomas, G. Tiao, M.T. Tusie-Luna, B. Weisburd, H.H. Won, D. Yu, D.M. Altshuler, D. Ardissino, M. Boehnke, J. Danesh, S. Donnelly, R. Elosua, J.C. Florez, S.B. Gabriel, G. Getz, S.J. Glatt, C.M. Hultman, S. Kathiresan, M. Laakso, S. McCarroll, M.I. McCarthy, D. McGovern, R. McPherson, B.M. Neale, A. Palotie, S.M. Purcell, D. Saleheen, J.M. Scharf, P. Sklar, P.F. Sullivan, J. Tuomilehto, M.T. Tsuang, H.C. Watkins, J.G. Wilson, M.J. Daly, D.G. MacArthur, C. Exome Aggregation, Analysis of protein-coding genetic variation in 60,706 humans, Nature 536(7616) (2016) 285–91.

[40] J.B. Holmes, E. Moyer, L. Phan, D. Maglott, B. Kattman, SPDI: Data Model for Variants and Applications at NCBI, Bioinformatics (2019).

[41] H. Obinata, S. Gutkind, J. Stitham, T. Okuno, T. Yokomizo, J. Hwa, T. Hla, Individual variation of human S1P(1) coding sequence leads to heterogeneity in receptor function and drug interactions, J Lipid Res 55(12) (2014) 2665–75.

[42] A.L. Parrill, D. Wang, D.L. Bautista, J.R. Van Brocklyn, Z. Lorincz, D.J. Fischer, D.L. Baker, K. Liliom, S. Spiegel, G. Tigyi, Identification of Edg1 receptor residues that recognize sphingosine 1-phosphate, J Biol Chem 275(50) (2000) 39379–84.

[43] Exome Variant Server, NHLBI GO Exome Sequencing Project (ESP), Seattle, WA (URL: http://evs.gs.washington.edu/EVS/)

[44] Y. Zhang, Y. Huang, A. Cantalupo, P.S. Azevedo, M. Siragusa, J. Bielawski, F.J. Giordano, A. Di Lorenzo, Endothelial Nogo-B regulates sphingolipid biosynthesis to promote pathological cardiac hypertrophy during chronic pressure overload, JCI Insight 1(5) (2016).

