## Supplemental Data for "Deletion of Sphingosine 1-Phosphate Receptor 1 in cardiomyocytes during development leads to abnormal ventricular conduction and fibrosis"

**Supplemental Movie 1:** Movie file corresponding to Figure 1B.
**Supplemental Movie 2:** Movie file corresponding to Figure 1C.
**Supplemental Movie 3:** Movie file corresponding to Figure 1D.
**Supplemental Movie 4:** Movie file corresponding to Figure 1E.
**Supplemental Movie 5:** Movie file corresponding to Figure 1F.
**Supplemental Movie 6:** Movie file corresponding to Figure 1G.

| ***Mlc2a* locus** | ***Sphk1* locus** | ***Sphk2* locus** | **Weaned mice** |
|---|---|---|---|
| +/+ | f/+ | -/- | 14 (9) |
| | f/- | -/- | 9 (9) |
| Cre/+ | f/+ | -/- | 5 (9) |
| | f/- | -/- | 8 (9) |

**Supplemental Table 1.** Generation of S1P ligand in cardiomyocytes is not required for survival. Genotypes are shown for pups from *Mlc2a*$^{Cre/+}$; *Sphk1*$^{+/-}$; *Sphk2*$^{-/-}$ mice crossed with *Sphk1*$^{f/f}$; *Sphk2*$^{-/-}$ mice. The observed number of weaned mice for each genotype is followed by the expected number of weaned mice in parentheses. All genotypes were represented at the expected numbers ($\chi^2$ 4.556, p = 0.1979).

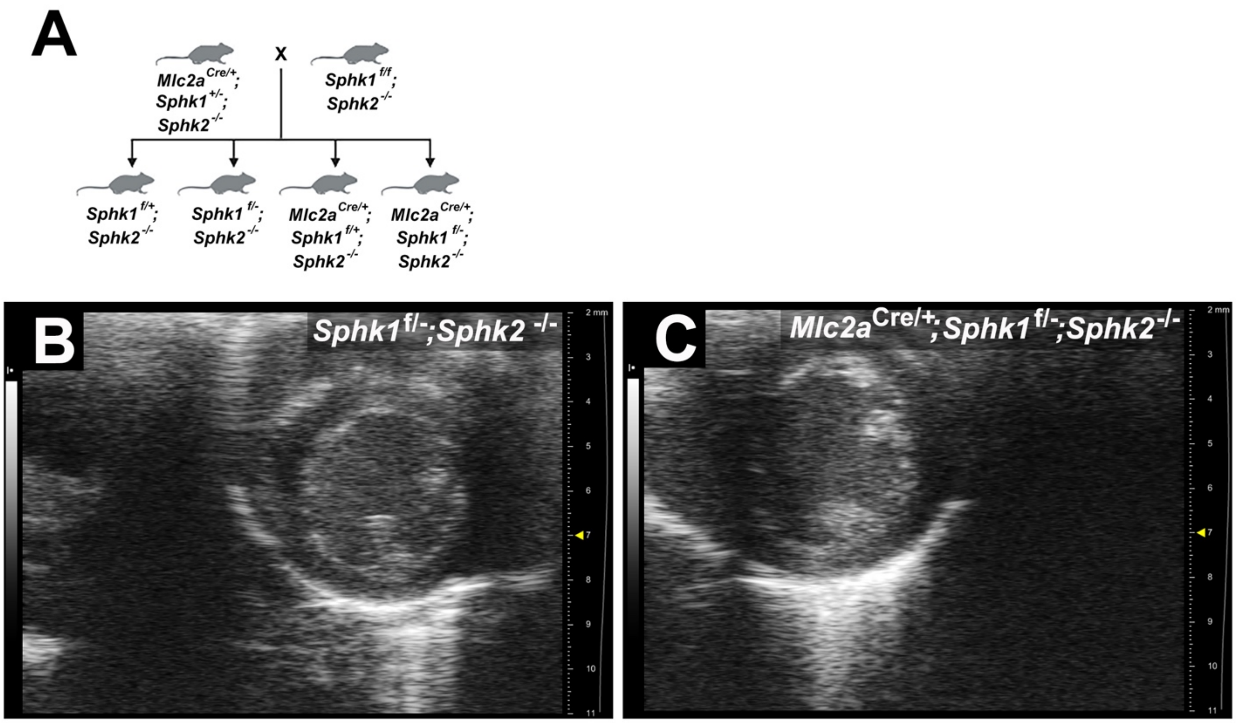


**Supplemental Figure 1. Generation of S1P ligand in cardiomyocytes is not required for normal heart development.** A) Breeding strategy. B) Parasternal short axis view from a *Sphk1*$^{f/-}$; *Sphk2*$^{-/-}$ control mouse. C) Parasternal short axis view from an *Mlc2a*$^{Cre/+}$; *Sphk1*$^{f/-}$; *Sphk2*$^{-/-}$ littermate mutant mouse. Note the absence of hypertrabeculated myocardium in the *Mlc2a*$^{Cre/+}$; *Sphk1*$^{f/-}$; *Sphk2*$^{-/-}$ mutant heart.

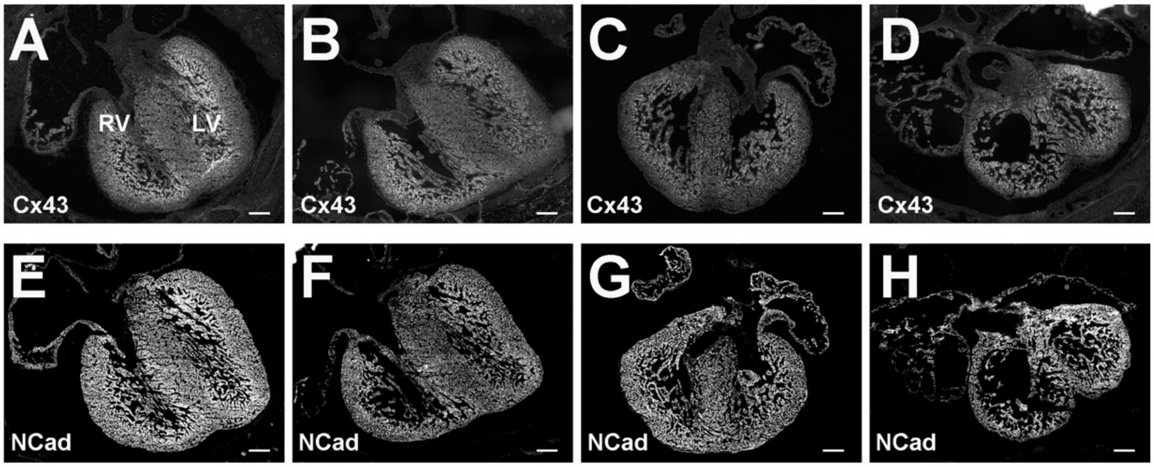


**Supplemental Figure 2. Connexin 43 (Cx43) and N-Cadherin (NCad) immunostaining in hearts from embryos collected at 15.5 dpc.** Representative images from n = 3 per genotype are shown. A, E) *S1pr1*$^{+/+}$ embryonic heart. B, F) *S1pr1*$^{+/-}$ embryonic heart. C, G) *Mlc2a*$^{Cre/+;}$ *S1pr1*$^{f/+}$ embryonic heart. D, H) *Mlc2a*$^{Cre/+;}$ *S1pr1*$^{f/-}$ mutant embryonic heart. No significant differences were noted among the four genotypes. LV, left ventricle. RV, right ventricle. Scale bar, 200 μm.

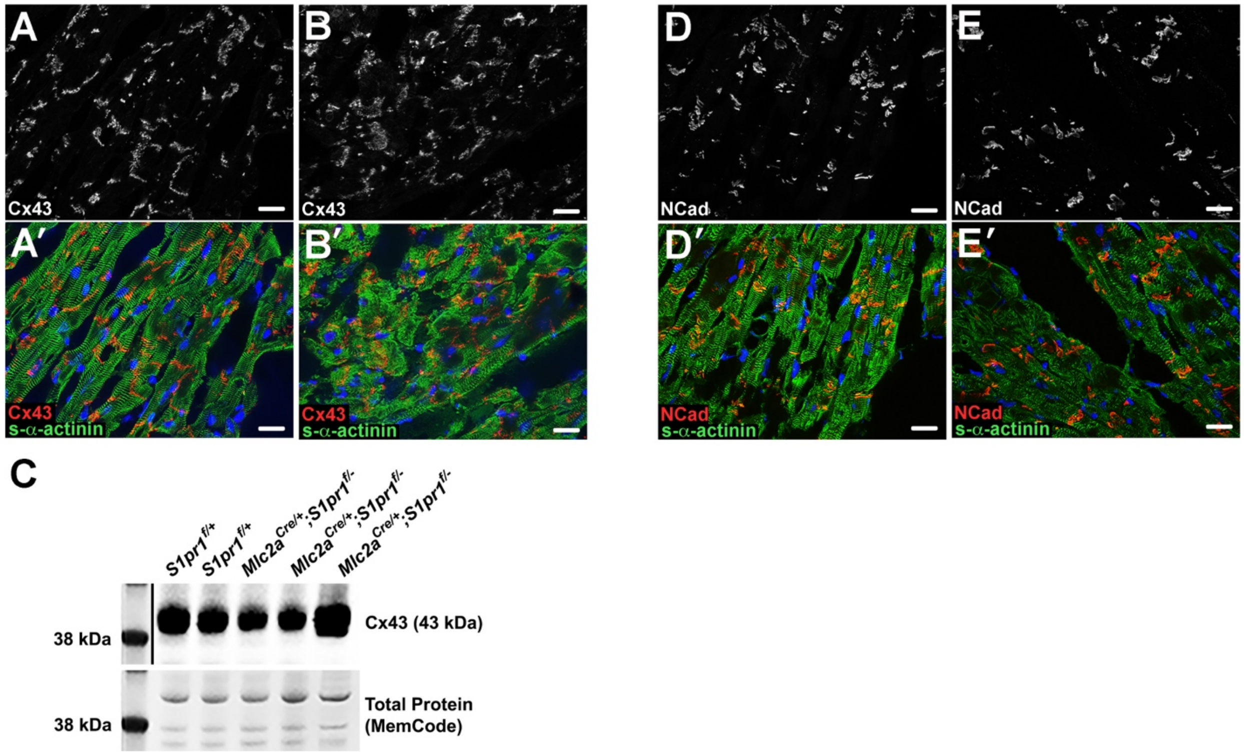


**Supplemental Figure 3. Normal expression of connexin 43 (Cx43) and N-Cadherin (NCad) in mice with embryonic cardiomyocyte *S1pr1* deletion.** Adult hearts were perfused with cardioplegia solution, snap frozen, cryosectioned, and immunostained as indicated. Representative images from n = 3-6 mice per genotype are shown.

A*)* *S1pr1*$^{f/+}$ control heart immunostained for Cx43.

A') Merged immunostaining for s-α-actinin (myofibril marker) and Cx43 in the *S1pr1*$^{f/+}$ control section shown in panel A.

B) *Mlc2a*$^{Cre/+;}$ *S1pr1*$^{f/-}$ mutant heart immunostained Cx43.

B') Merged immunostaining for s-α-actinin and Cx43 in the *S1pr1*$^{f/+}$ *Mlc2a*$^{Cre/+;}$ *S1pr1*$^{f/-}$ mutant section shown in panel B.

C) Immunoblot of lysates from whole left ventricle and right ventricle show no consistent difference in Cx43 levels between *S1pr1*$^{f/+}$ control and *Mlc2a*$^{Cre/+;}$ *S1pr1*$^{f/-}$ mutant hearts.

D) *S1pr1*$^{f/+}$ control heart immunostained for NCad.

D') Merged immunostaining for s-α-actinin and NCad in the *S1pr1*$^{f/+}$ control section shown in panel D.

E) *Mlc2a*$^{Cre/+;}$ *S1pr1*$^{f/-}$ mutant heart immunostained for NCad.

E') Merged immunostaining for s-α-actinin and NCad in the *Mlc2a*$^{Cre/+;}$ *S1pr1*$^{f/-}$ mutant section shown in panel E.

No significant differences were noted among the genotypes. Scale bar, 20 μm.
